# *SlTDP1* Is Required To Specify Tapetum Identity And For The Regulation Of Redox Homeostasis In Tomato Anthers

**DOI:** 10.1101/2021.12.03.471129

**Authors:** Blanca Salazar-Sarasua, María Jesús López-Martín, Edelín Roque, Rim Hamza, Luis Antonio Cañas, José Pío Beltrán, Concepción Gómez-Mena

## Abstract

The tapetum is a specialized layer of cells within the anther adjacent to the sporogenic tissue. During its short life, it provides nutrients, molecules and materials to the pollen mother cells and microsporocytes being essential during callose degradation and pollen wall formation. However, the acquisition of tapetal cell identity in tomato plants is a process still poorly understood. We report here the identification and characterization of *SlTPD1* (*Solanum lycopersicum TPD1*), a gene specifically required for pollen development in tomato plants. Gene editing was used to generate loss-of-function *Sltpd1* mutants that showed absence of tapetal tissue. In these plants, sporogenous cells developed but failed to complete meiosis resulting in complete male sterility. Transcriptomic analysis conducted in wild-type and mutant anthers at an early stage revealed the down regulation of a set of genes related to redox homeostasis. Indeed, *Sltpd1* anthers showed a reduction of reactive oxygen species (ROS) accumulation at early stages and altered activity of ROS scavenging enzymes. The obtained results highlight the importance of ROS homeostasis in the interaction between the tapetum and the sporogenous tissue in tomato plants.

**One sentence summary:** The small protein SlTPD1 is required for tapetum formation in tomato, highlighting the role of this tissue in the regulation of redox homeostasis during male gametogenesis.

## INTRODUCTION

Sexual reproduction in both animals and plants requires the formation of haploid gametes in a complex and highly regulated process. Quite unlike animals, in flowering plants the gametes are produced post embryonically within specialized organs, the ovary and the anther. The female gametophyte (embryo sac) is produced from a germline originated in the ovules inside the ovary, while male gametophytes (pollen) originate inside the anther. The formation of gametes in plants occurs late in development and does not depend on meristems but on cell-to-cell communication or tissue interactions.

The anther shows a relatively simple morphological structure and high accessibility being the object of numerous studies on the sexual reproduction of plants. Shortly after anther primordia initiation, several somatic and germinal cells originate. Typically, the primordia contains three cells layers (L1-L3) that will originate the external epidermis, the archesporial cells and the inner vascular and connective tissue (Gómez et al., 2015; Åstrand et al., 2021). Archesporial cells further differentiate into three additional layers of somatic tissue: the endothecium, the middle layer and the tapetum, and a layer of microsporocytes (pollen mother cells, PMC). The tapetum layer, adjacent to the developing microsporocytes has a central role during pollen development and its premature or delayed degradation results in pollen abortion and male sterility (Liu et al., 2018; Bai et al., 2019).

Most of the genetic information on male gametogenesis was obtained in the model plant *Arabidopsis thaliana* and two monocot crops, rice and maize (Chang *et al*., 2011; van der Linde & Walbot, 2019). In Arabidopsis, tapetal cell formation requires the joined action of EMS1/EXS (EXCESS MICROSPOROCYTES1/EXTRA SPOROGENOUS

CELLS) a putative Leucin-rich repeat (LRR) receptor kinase (Canales et al., 2002; Zhao et al., 2002) and its ligand the small peptide TPD1 (TAPETUM DETERMINANT1)(Yang et al., 2003; Huang et al., 2016b). In rice, a similar receptor/ligand complex is encoded by the *MSP1* (*MULTIPLE SPOROCYTE1)* and *TDL1A* genes (Nonomura et al., 2003; Zhao et al., 2008) while in maize a *TPD1* homolog, *MAC1(MULTIPLE ARCHESPORIAL CELLS1*)/*MIL2* gene (Hong et al., 2012a; Wang et al., 2012), was identified. Downstream of this complex, several genes such as *BRI1 EMS SUPPRESSOR* (*BES1*), *DYT1* (*DYSFUNCTIONAL TAPETUM1*), *DEFECTIVE IN TAPETAL DEVELOPMENT AND FUNCTION1* (*TDF1*) and *MYB33/65* are required for early tapetal development and function in Arabidopsis (Millar and Gubler, 2005; Zhu et al., 2008; Gu et al., 2014; Chen et al., 2019). At late stages, *MALE STERILITY1* (*MS1*) and *AMS* regulate pollen formation and maturation (Ito and Shinozaki, 2002; Sorensen et al., 2003). Despite small differences, extensive research in Arabidopsis and rice suggests that the genetic pathway controlling tapetum development is highly conserved in plants (Wilson and Zhang, 2009; Zhang and Yang, 2014; Lei and Liu, 2020).

In tomato, male sterility is a desirable trait to be used in hybrid seed production and cross breeding programs. Over 50 male sterile mutants were isolated more than two decades ago (Gorman and McCormick, 1997) and still only a limited amount of genes involved in male gametogenesis have been identified. Mutations in the tomato *SPOROCYTELESS/NOOZLE* orthologue prevent the formation of both male and female sporocytes and the plants are fully sterile (Hao *et al*., 2017; Rojas-Gracia *et al*., 2017). Downstream of this gene, *Ms10^35^* gene (*DYT1* homolog) encodes a bHLH transcription factor specifically expressed in tapetal tissue and meiocytes (Jeong et al., 2014). Another bHLH protein (Solyc01g081100) has been proposed as the best candidate to encode the tomato *Ms32* gene (Liu et al., 2019). *Solyc01g081100* gene is a homolog of the Arabidopsis *bHLH10/89/90* gene that together with DYT1-MYB35 form a regulatory module to regulate tapetum and pollen development (Cui et al., 2016).

In this work, we identified a gene that is specifically required for the specification of the tapetal cells in tomato. The gene corresponds to the tomato homolog of the *TPD1* Arabidopsis gene, and was named *SlTPD1* (*Solanum lycopersicum TPD1*). We obtained mutant plants by CRISPR/Cas9 technology that showed a male sterile phenotype associated with the absence of tapetal tissue. We studied the cytological and molecular changes of the anther and in particular the effect in sporogenous cell development in the mutant plants. Our results provide evidence for a regulatory role of the tapetum in the progression of male gametogenesis through the modulation of redox homeostasis.

## RESULTS

### Identification of the Solanum lycopersicum SlTPD1 gene

Following a gene homologue strategy, we selected a gene candidate to be involved in tapetum development in tomato. *TPD1* (TAPETUM DETERMINANT1) (Ya*ng et al*., 2003) was used as a bait in the Plant Comparative platform Phytozome (Goodstein et al., 2012) (https://phytozome-next.jgi.doe.gov/) and two homologous were identified (*Solyc03g097530* and *Solyc11g012650*). The expression of these genes was analyzed in vegetative tissues (leaves) and flower buds using qPCR. The results showed that *Solyc11g012650* was preferentially expressed in leaves while *Solyc03g097530* was expressed in developing flowers, reaching the highest level in flower at anthesis (Figure **S1**). Phylogenetic analyses were performed using a list of homologue genes from different plant species obtained in a BLAST search using *TPD1* gene (*At4g24972*) as a bait. These sequences also included the Arabidopsis closest homolog *At1g32583* and the rice orthologue *OsTDL1A* (Zhao et al., 2008). In the phylogenetic tree, *Solyc03g097530* grouped with *TPD1* and related TPD1-like homologues from Solanaceae (Figure **1A**).

**Figure 1.**
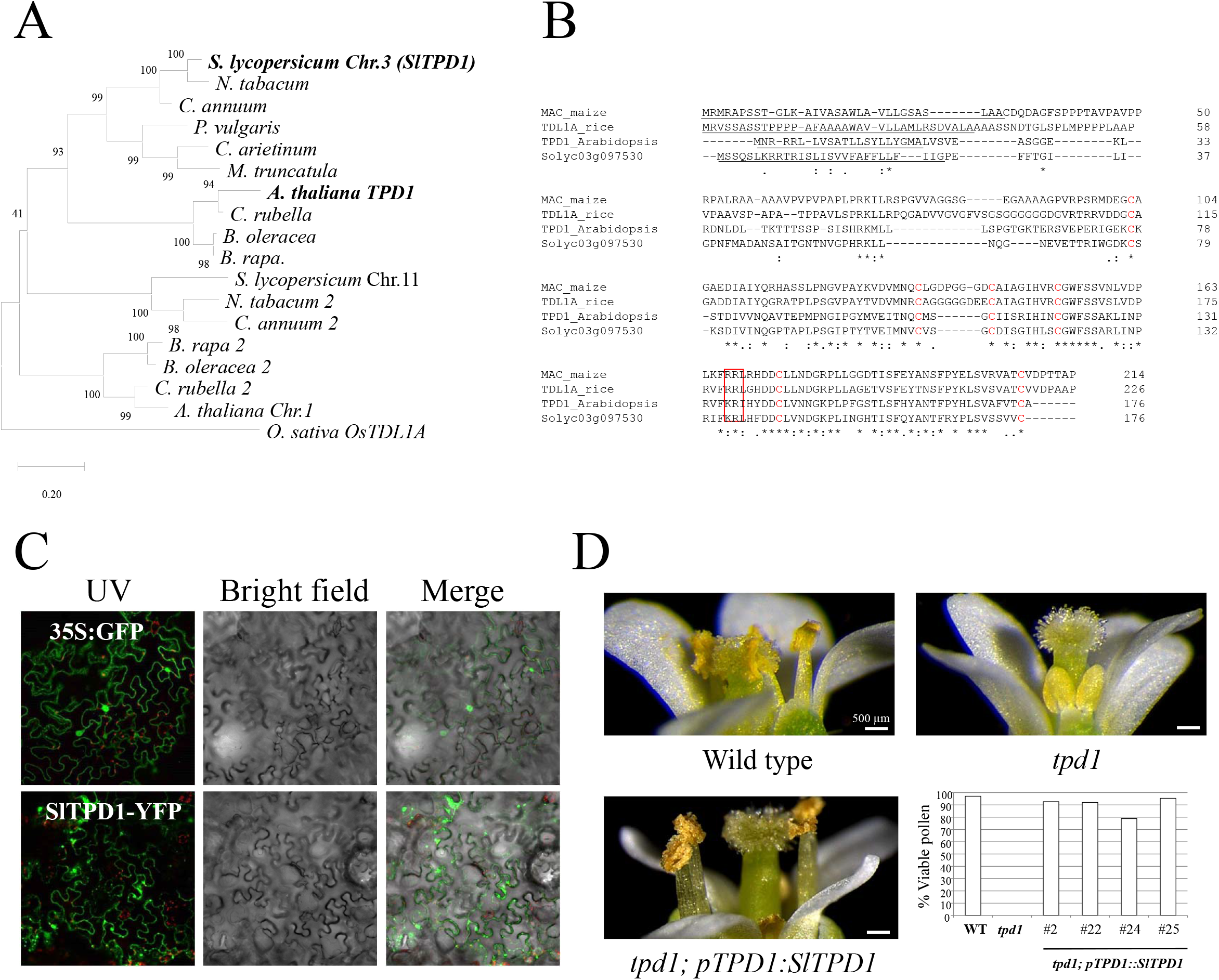
*Solyc03g097530* (*SlTPD1)* encodes the ortholog of *TPD1* in tomato. A, Unrooted neighbor-joining tree of TPD1-like proteins. The numbers next to the internal nodes are bootstrap values from 1000 pseudo-replicates. B, Amino acid sequence alignment between the Arabidopsis and tomato gene homologs. The putative signal peptides are underlined, the six conserved cysteine residues are in bold and the potential dibasic cleavage site is highlighted with a red square. C, Subcellular localization of SlTPD1 protein in *Nicotiana benthamiana* leaves as observed by confocal microscopy. D, Complementation of the male sterile floral phenotype of the Arabidopsis *tpd1* mutant using *SlTPD1* gene. All the *tpd1; pTPD1:SlTPD1* plants showed viable pollen. Scale bars in (D) correspond to 500 µm.

Solyc03g097530 protein sequence (176aa) was aligned with Arabidopsis TPD1 and two protein homologs functionally characterized: TDL1A from rice (Zhao *et al*, 2008) and MAC1 from maize (Wang et al., 2012). The proteins showed high amino acid identity mainly in the C-terminal region with six highly conserved cysteine residues and a putative dibasic cleavage site (Figure **1B**). In addition, SlTPD1 protein and homologs contain a predicted signal peptide at their N-terminal regions (Figure **1B**; underlined). The subcellular location of the protein was determined by fusing the Yellow Fluorescent protein (YFP) to the C terminal end of SlTPD1 and transiently expressed in *Nicotiana benthamiana* leaves. The control protein (35S:GFP) exhibited both cytoplasmic and nuclear localization (Figure **1C**) while SlTPD1-YFP protein was localized in proximity of the plasma membrane where it formed small dots, and in the cytosol as large aggregates (Figure **1C**). This result suggests that SlTPD1 protein could be secreted to the extracellular space.

Gene orthology and local microsynteny or collinearity was inferred by *in silico* analyses. The use of the Gene Orthology View in the PLAZA platform (Van Bel et al., 2018) confirmed that *Solyc03g097530* is the best orthologous candidate for *TPD1* in tomato (Figure **S2A**). We then looked for microsynteny in the flanking regions where the two genes are located and found collinearity between these two regions (Figure **S2B**). Therefore, we considered *Solyc03g097530* the strongest candidate to be the *TPD1* orthologous in tomato and renamed it *SlTPD1* (*Solanum lycopersicum TPD1*).

To evaluate the conservation of *SlTPD1* function during the development of the anther we designed an experiment to complement a loss-of-function *tpd1* mutant. We used a mutant line (N843482, *tpd1* mutant) that contains a T-DNA insertion in the *TPD1* gene. Mutant plants were indistinguishable from the wild type except for the anthers that did not produce pollen grains (Figure **1C**). To complement the mutant phenotype, we generated a genetic construct by fusing 2.7 kb of the promoter region of Arabidopsis *TPD1* and the coding sequence of *SlTPD1* that was used to genetically transform heterozygous *tpd1* plants. We obtained 32 independent transformants and four of them were homozygous for the mutation. These four plants produced viable pollen (Figure **1D**) and seeds and therefore, were fertile, demonstrating the ability of SlTPD1 protein to replace TPD1 function.

### Expression of *SlTPD1* during tomato plant development

The expression of *SlTPD1* was analyzed in different plant tissues including seedlings (apical and basal regions), leaves and developing flowers using qPCR. The gene was expressed in all the tissues analyzed reaching the highest level in flowers at anthesis (Figure **2A**). The spatial and temporal pattern of expression of *SlTPD1* was evaluated during flower development using *in situ* hybridization (Figure **2B-G**). *SlTPD1* RNA was not detectable in inflorescence meristems and flower buds before anther primordium differentiated (Figure **2B**). Expression was first detected at floral stage 6 at the internal layers of the developing anther that will generate the sporogenous tissue (Figure **2C****, D**). Later, at the tetrad stage, *SlTPD1* transcript was localized at the tapetum and the microspores still surrounded by the callose wall. The expression of the gene continues during the following floral stages in the tapetal cells that gradually disintegrated and in the pollen grains (Figure **2E-G**). On the ovary, we detected transient expression in ovule primordia of flower at stage 8 (Figure **S3**).

**Figure 2.**
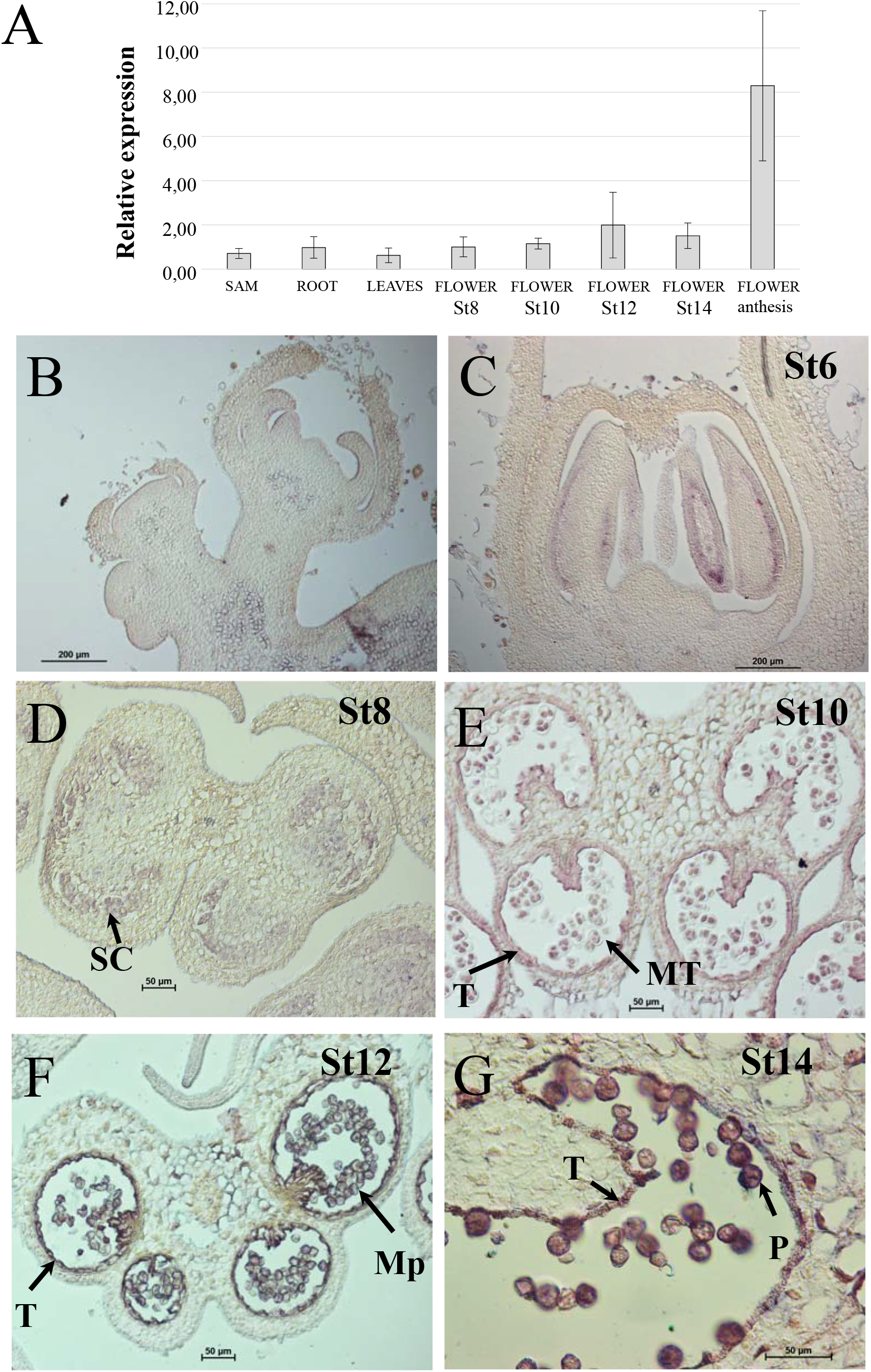
Expression of *SlTPD1* during plant development. A, Relative expression of *SlTPD1* in different plant tissues analysed by qPCR. Data were normalized to the expression of *ACT10* gene and correspond to the mean (±SD) of three biological replicates. B-G, Localization of *SlTPD1* transcript by *in situ* hybridization on reproductive meristems and developing flowers. St6: floral stage 6; St8: floral stage 8; St10: floral stage 10; St12: floral stage 12; St14: floral stage 14. SC: Sporogenous cells; MT: microspore tetrads; T: Tapetum; Mp: microspores; P: mature polen.

### *Sltpd1* mutants are male sterile and developed parthenocarpic fruits

To study the function of *SlTPD1*, tomato lines with mutations targeted to the third exon were generated using CRISPR/Cas9 (Figure **3A**). Among the T0 generation, we selected six diploid plants that showed percentages of edition over eighty and that mostly contained biallelic mutations (Figure **S4A**). All the plants showed complete male sterility and developed seedless (parthenocarpic) fruits. Histological sections of the mature anthers revealed collapsed locules containing a dense debris but they did not contain viable pollen (Figure **S4B**). In these plants, we observed a strong correlation between male sterility and the development of parthenocarpic fruits (Figure **S4C**).

**Figure 3.**
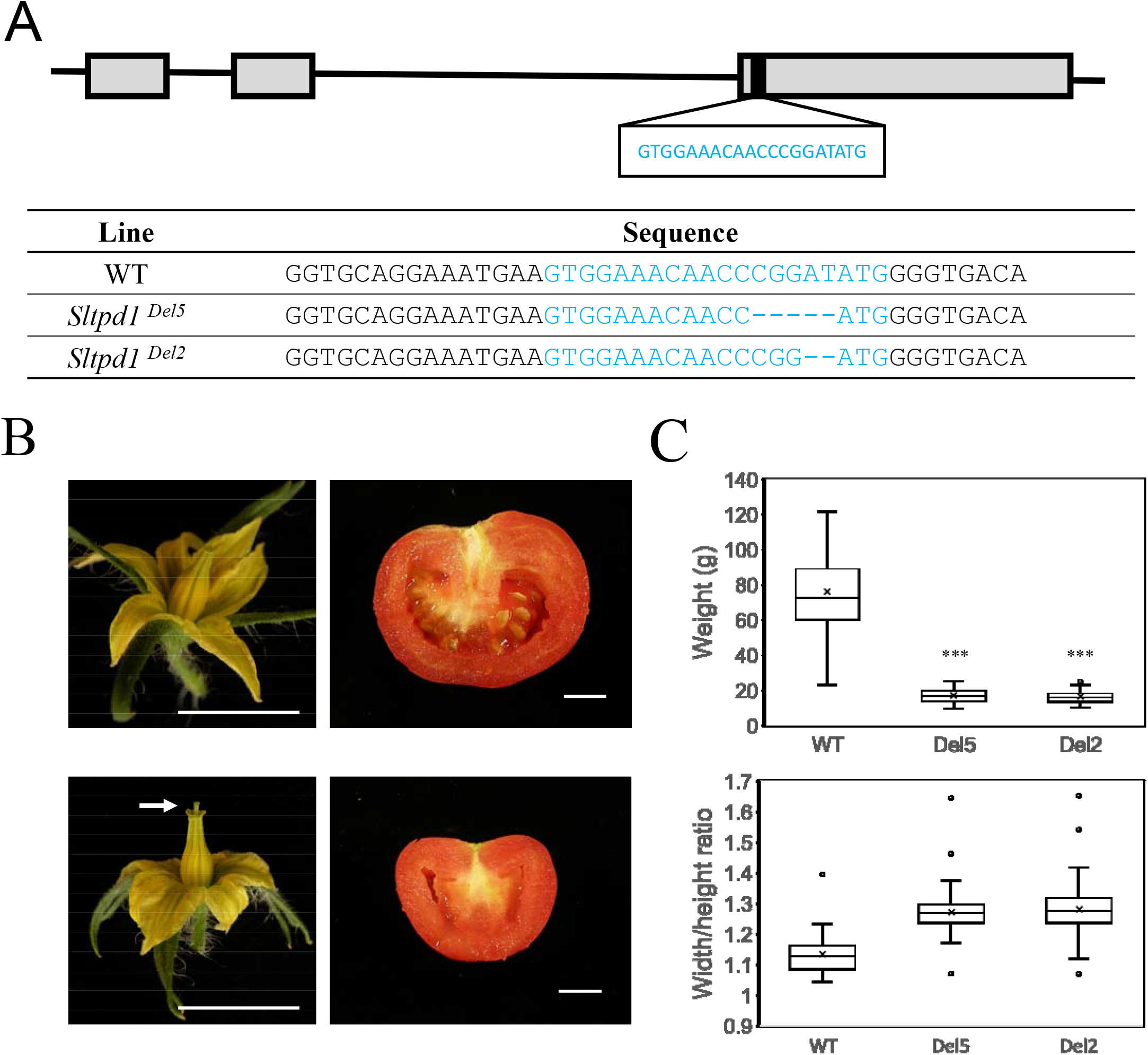
Mutations in *SlTPD1* result in empty anthers and seedless fruits. A, Guide RNA (in blue) was targeted to the third exon of *SlTPD1*. B, Wild-type and *Sltpd1* flowers and opened fruits. *Sltpd1* plants showed flowers with protruding pistils (arrow) and seedless fruits. Scale bar: 0.5 cm. C, Fruit weight and shape (width/height ration) of the wild type, *Sltpd1* ^Del5^ and *Sltpd1* ^Del2^ plants (n ≥ 40 ± SD).

F2 plants were obtained after pollination with wild-type pollen and stable single-mutation lines were obtained. Two mutant lines (*Sltpd1^Del5^ and Sltpd1^Del2^*) containing deletions of 2 and 5 nucleotides respectively were chosen for further analysis (Figure **3A**). Mutant plants did not show morphological defects during vegetative development. However, after flower opening, we observed a small reduction in the stamen length, absence of pollen and slight protrusion of the pistil (Figure **3A**). For many crops, including tomato, the production of hybrids is an efficient way to increase plant production and improve their resistant to diseases and its performance under suboptimal environmental conditions (Labroo et al., 2021). Male-sterile tomato plants with protruding stiles, such as *Sltpd1* mutants, could be valuable parental lines for hybrid seed production. Despite male sterility, *Sltpd1* mutant plants produced seedless fruits that were smaller than those of wild-type plants with a decrease of about 70% in weight (Figure **3C**). Fruit shape, quantified as a width/height ratio, was not altered in *Sltpd1* mutants (Figure **3C**). When fertilized with wild-type pollen, the plants developed seeded fruits of a normal size.

### Male gametogenesis fails to be completed in *Sltpd1* mutants

To elucidate the biological function of *SlTPD1* during male gametogenesis, we compared the development of anthers from the wild type and *Sltpd1* mutants. In wild-type tomato anthers, cells from the L2 layer differentiate into archesporial cells that undergo periclinal divisions (parallel to the epidermis) (Figure **4A**). In *Sltpd1* mutant, anther development was slightly different to the wild type showing cells with squared rather than rectangular shape and reduced number of periclinal divisions (Figure **4D**). From stage 8, we observed clear differences between the two genotypes. While epidermis, endothecium and middle cell layers were formed, tapetum was not present in the mutant and sporogenous cells seemed more abundant and disorganized compared to the wild type (Figure **4E**). At stage 10, wild-type microsporocytes completed meiosis and formed tetrads surrounded by callose and separated from the adjacent cell layers (Figure **4C**). Eventually, callose was degraded, releasing the microspores that continued to develop into mature pollen grains during floral stages 12 to 16 (Figure **4G-I**). Simultaneously, the tapetum started to degrade and was not visible by stage 16 (Figure **4I**). In the mutant anthers, microsporogenous cells continued to divide and enlarged in size (Figure **4F**). After extra rounds of divisions, the cells occupied the complete cavity of the locule (Figure **4J**). Cell counting showed that by floral stage 8 the number of sporogenous cells in *Sltpd1* anther locules roughly doubled that of wild-type anthers (24.0 ± 5.8 versus 49.0 ± 4.7 cells/locule section). At stage 10, sporogenous cell number further increased (66.1 ± 11.3 cells/locule section) and cells seemed to have initiated meiosis but failed to complete it (Figure **4J**). Finally, cells degenerated causing the collapse of the anther locules and the deposition of a dense cell debris (Figure **4L****, L**).

**Figure 4.**
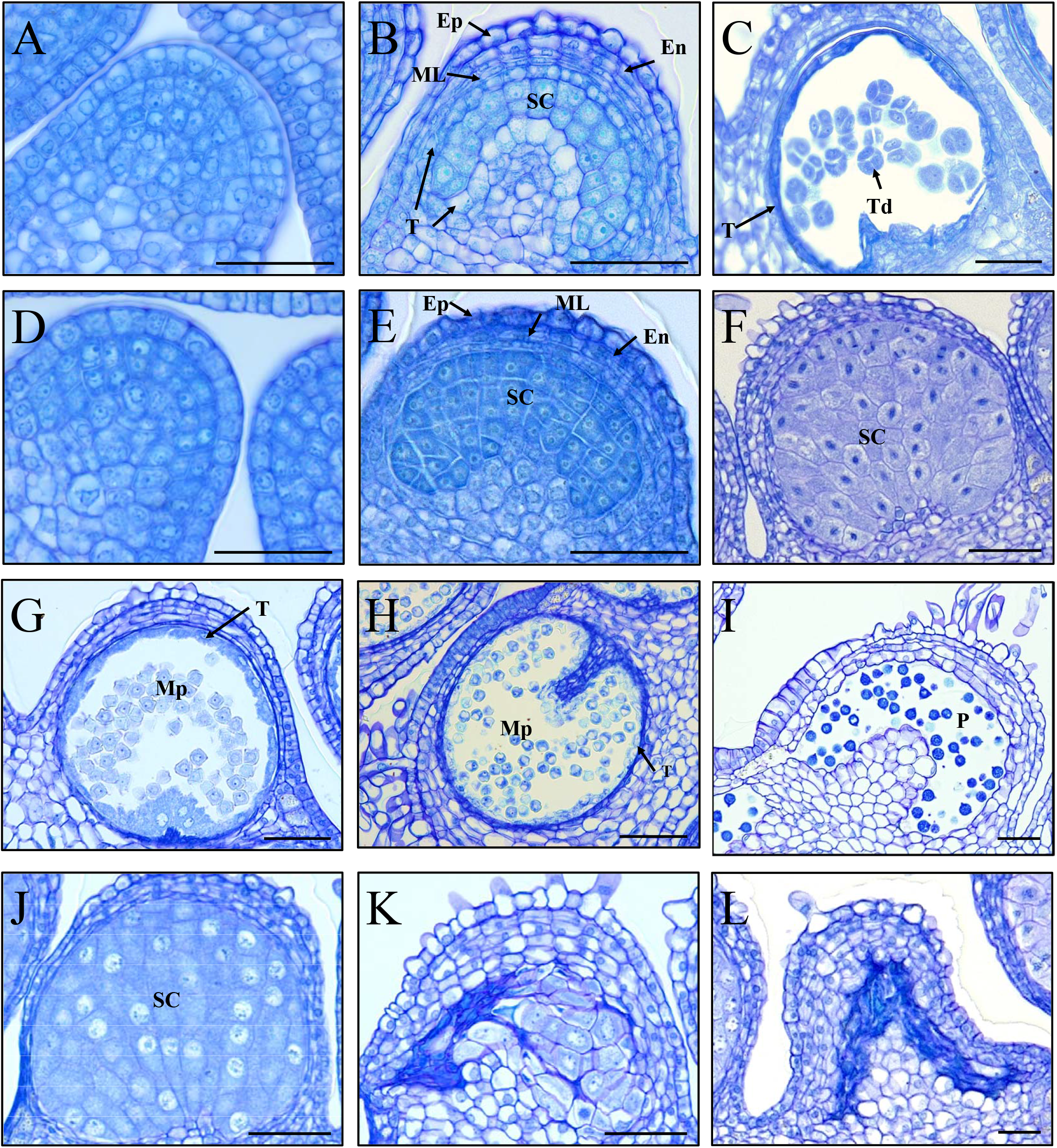
Pollen development is impaired in *Sltpd1* mutants. A-L, Histological sections of anther from the wild type (a-c and g-i) and *Sltpd1* mutant (D-F and J-L) at different developmental stages. Transveral section of anthers from floral stage 6 (A) and (D), floral stage 8 (B) and (E), floral stage 10 (C) and (F), floral stage 12 (G) and (J), floral stage 14 (H) and (K) and floral stage 16 (I and L). Floral stages have been named according to Brukhin et al 2003. Scale bar: 50 µm. Ep: epidermis; En: endotecium; ML: middle layers; SC: sporogenous cells; T: tapetum; Td: tetrads; Mp: microspores; P: mature pollen.

We performed *in situ* hybridization essays using probes for the tapetum-specific *TomA5B* (*Solyc01g086830*) gene (Aguirre and Smith, 1993) and for the tomato homologue (*Solyc04g008070*) of the meiosis marker *SOLO DANCER* (*SDS*) gene (Azumi et al., 2002). In the wild type anther, *TomA5B* probe strongly hybridized with the tapetal cells at floral stages 8 and 10 (Figure **5A****, B**). The signal decreased dramatically by stage 12 when tapetum degeneration starts (Figure **5A**). In the *Sltpd1* mutant anthers no signal was obtained in any of the floral stages analyzed (Figure **5D-F**) confirming the complete absence of tapetum in the mutant plants. In the case of *SlSDS* probe, the hybridization signal was first observed in the wild type at floral stage 8 overlapping with meiosis initiation (Figure **5B**). A similar result was obtained in the mutant anther detecting the expression of the gene at floral stage 8 (Figure **5B**). These results indicate that meiosis was initiated in the mutant anthers although it failed to progress to the tetrad stage.

**Figure 5.**
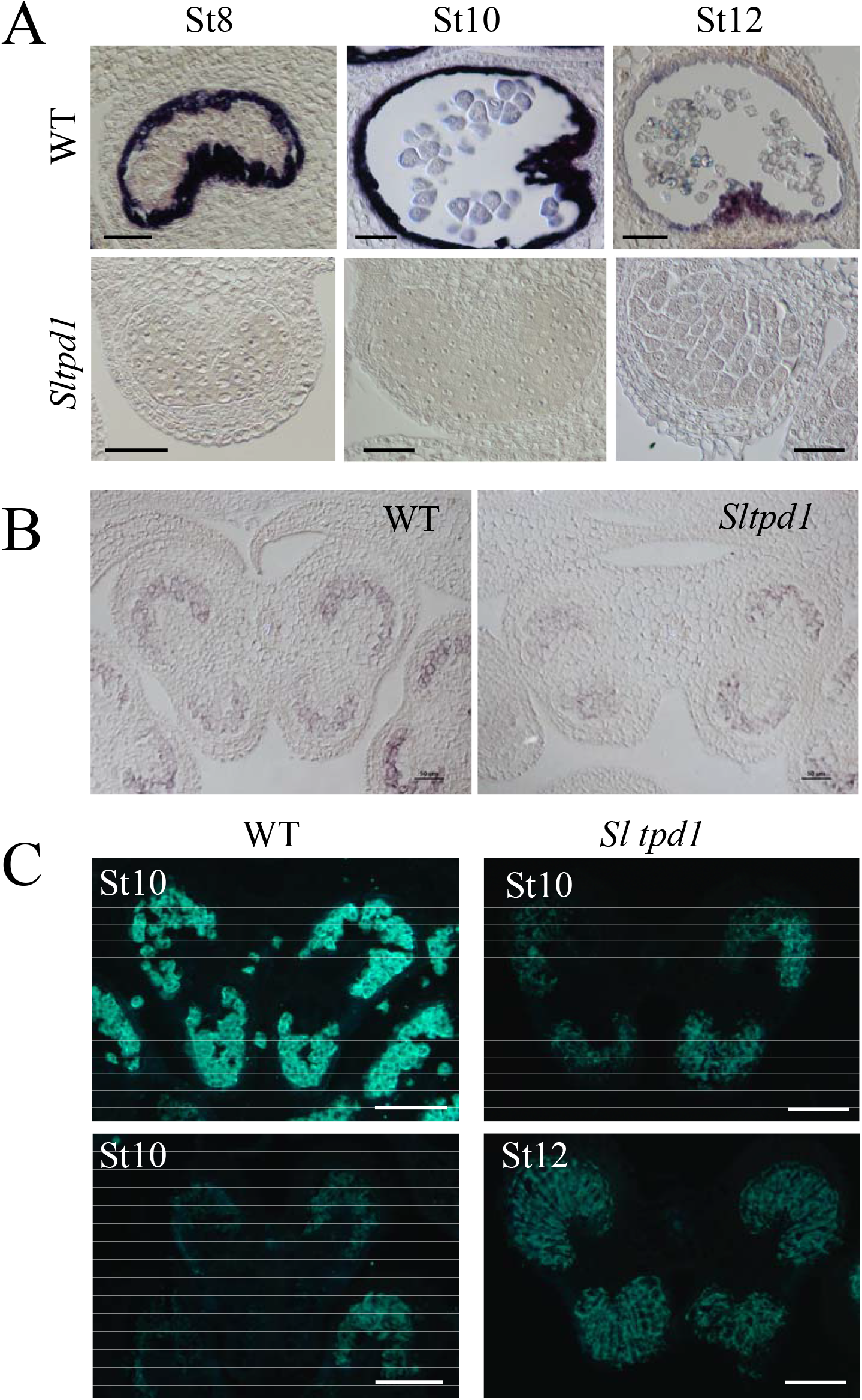
*Sltpd1* mutant anthers specifically lack tapetal cells. A, *In situ* hybridization of the tapetum marker *TomA5B* in wild type and *Sltpd1* anthers. B, *In situ* hybridization of the meiosis marker *SlSDS* in wild type and *Sltpd1* anthers at floral stage 8. C, Callose deposition in anthers as observed by aniline blue staining of wild type and *Sltpd1.* St8: floral stage 8; St10: floral stage 10; St12: floral stage 12. Scale bar: 50 µm in (A) and (B); 100 µm in (C).

Callose deposition occurs around the sporogenous cells predating meiosis initiation and later between meiotic products (Jaffri and MacAlister, 2021). After meiosis completion, callose is quickly degraded after the release of callases (ß1-3 glucanases) by the tapetum. Using aniline blue staining the pattern of callose deposition and degradation was analyze in the mutant plants. In wild type anthers, the deposition of callose appears as an intense florescence signal around the tetrads that quickly disappears at the termination of meiosis (Figure **5C**). In the mutant plants, the accumulation of callose was observed as a diffuse signal surrounding the sporogenous cells and the fluorescence signal persisted in time until the collapse of the anther locule (Figure **5C**).

### Identification of global transcriptional changes associated to *SlTPD1* loss-of-function

To identify molecular and cellular components downstream of *SlTPD1* action, RNAseq analyses were performed using anthers from floral stage 8 (meiotic stage). Differential expressed genes (DEG) were selected using a Q-value >0.1 and p-value >0.05. From the selected genes (801), 519 correspond to down-regulated genes and 282 to up-regulated genes (Figure **6A** and Table **S1**).

**Figure 6.**
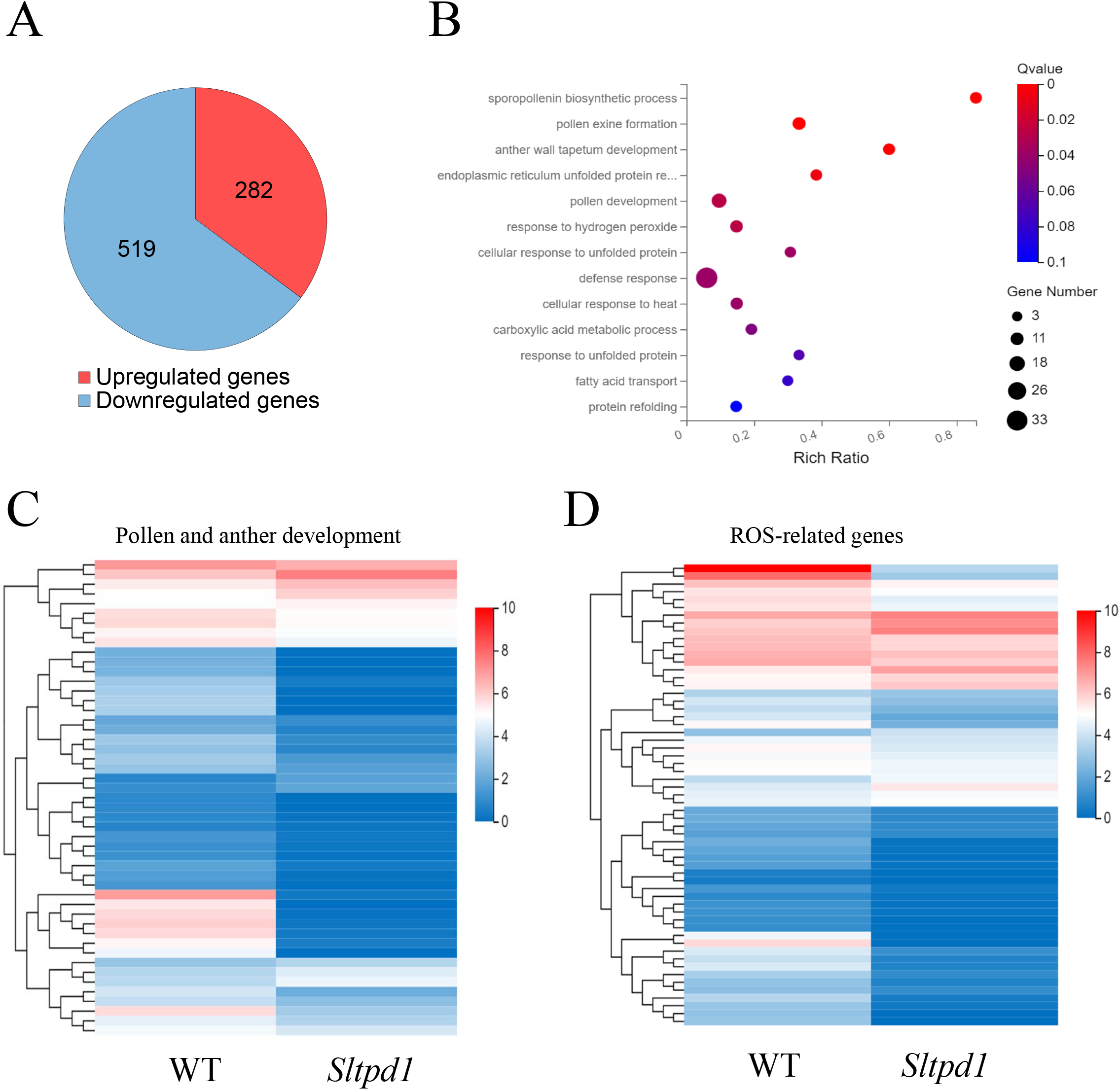
Global gene expression chages in the anthers of *Sltpd1* mutants at floral stage 8 in comparison with the wild type. A, Total number of DEGs between wild type and mutant anthers. B, GO biological process enrichment analysis. C, Expression heatmap of differentially expressed genes involved in pollen and anther development. D, Expression heatmap of differentially expressed ROS-related genes. Q-value < 0.05; p-value < 0.05.

**Table 1.**
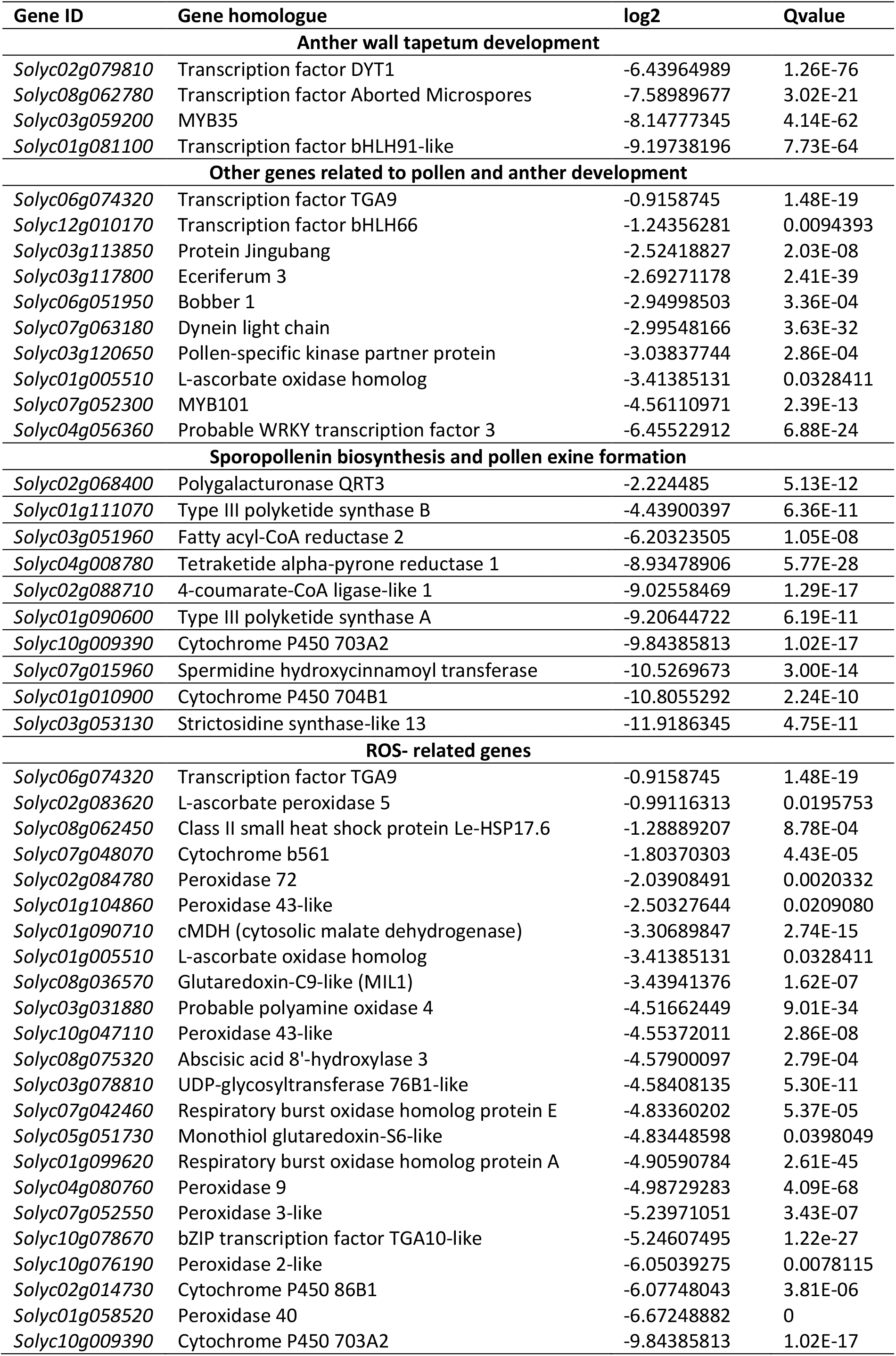
List of genes involved pollen development and reactive oxygen species (ROS) homeostasis that showed down regulation in the anthers of Sltpd1 mutants compared to the wild type at floral stage 8.

At early floral stage 8, Gene Onthology (GO) analyses revealed enrichment in genes related to pollen and tapetum development (five out thirteen categories that correspond to 49 genes; Figure **6B** and Table **S1**). Among these genes, homologs to *DYT1*, *AMS*, *MYB35* and *bHLH91* showed strong downregulation (Table **1**). Accordingly, in Arabidopsis DYT1 is required to activate the expression of *bHLH010/bHLH089/bHLH091* genes which in turn facilitate DYT1 nuclear location and promote *MYB35* expression (C*ui et al*., 2016). In addition, we detected strong downregulation of genes required during late stages of pollen development including a polygalacturonase homolog of the Arabidopsis *QRT3* gene involved in microspore separation (Rhee et al., 2003) and a fatty acid-CoA reductase (Table **1**).

When looking at signaling pathways, an important group of redox related genes was observed grouped under “cellular response to hydrogen peroxide” and “defense response”. A specific expression heat-map analysis of redox-related genes revealed differential expression of seventy genes, of which fifty-three were downregulated and seventeen were upregulated (Figure **6C** and Table **S1**). Among the downregulated genes (Table **1**), we detected two *Respiratory burst oxidase homolog* (*Rboh*) genes (also known as NADPH oxidases), key enzymes that catalyze the formation of ROS in plants and a glutaredoxin (GRX) that shows homology with the *MIL1* gene from rice involved in microspore development (Hong *et al*., 2012b). Moreover, nine peroxidases are downregulated in the mutant anthers including homologues of the previously characterized PRX9 and PRX40 involved in pollen development in Arabidopsis (Jacobowitz et al., 2019). Peroxidases are multifunctional proteins that catalyze the oxidation of a variety of substrates by H_2_O_2_ and act as efficient components of the antioxidative system controlling ROS.

We analyzed the contribution of the genes involved in redox homeostasis to the development of the tomato anther. From the list of DEGs a subset of key ROS-related genes was selected, and its expression level were checked in anthers from different developmental stages (St6 to St20). The expression of two tomato *RBOH* genes (*SlRbohA/Solyc01g099620* and *SlRbohE/ Solyc06g075570*) was analyzed by qPCR. Besides, we analyzed the expression of *SlRBOH1*/*SlRbohG*, recently identified as a brassinosteroid (BR)-regulated gene involved in tapetal cell degeneration and pollen development (Yan *et al*., 2020). In *Sltpd1* mutant anthers we detected an important reduction in the expression level of *SlRbohA* and *SlRbohE* at early stages of anther development (Figure **7A****, B**). The expression levels of *SlRBOH1*/ *SlRbohG* did not significantly change during the floral stages analyzed (Figure **7C**).

**Figure 7.**
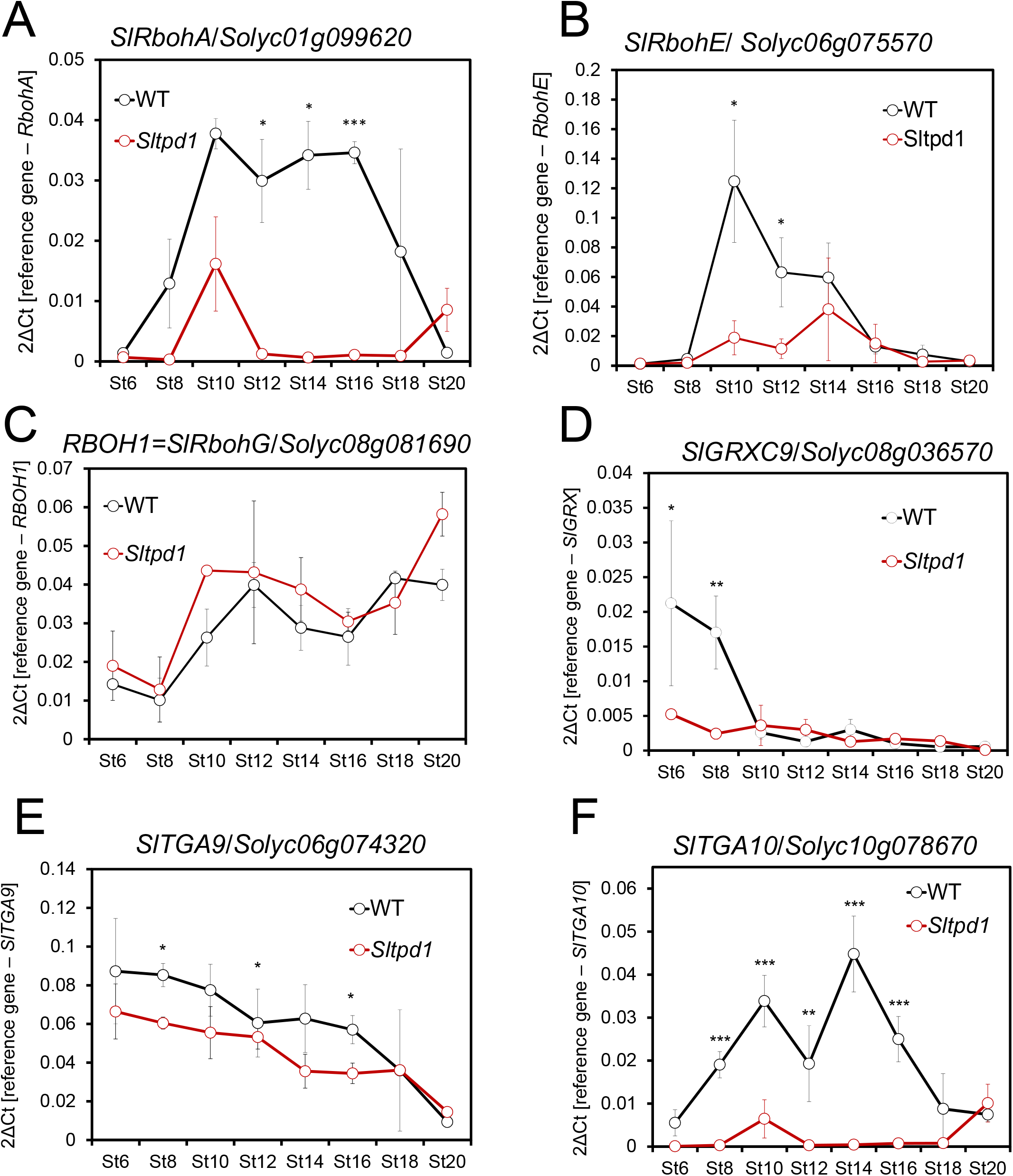
Expression pattern of genes involved in redox homeostais during anther development of the wild type and *Sltpd1* mutant plants. Quantitative RT-PCR of (A) *SlRbohA/Solyc01g099620* gene; (B) *SlRbohE/Solyc06g075570* gene; (C) *RBOH1SlRbohG/Solyc08g081690* gene; (D) *SlGRX9/Solyc08g036570* gene; (E) *SlTGA9/Solyc06g074320* gene and (F) *SlTGA10/Solyc10g078670* gene. Data correspond to three biological replicates ± SD. Statistical differences were inferred using a Mann-Whitney test. (*) = p<0.05, (**) = p<0.01, (***) = p<0.001.

The expression of the glutaredoxin-C9-like gene *SlGRXC9/Solyc08g036570* was analyzed and high levels of expression were detected in the wild type at the earliest stages analyzed (floral stage 6 and 8), while in the mutant samples the expression level was greatly reduced (Figure **7A**). The expression of two TGA-like transcription factors (*Solyc06g074320/SlTGA9* and *Solyc10g078670/SlTGA10*), downregulated in the RNA-seq, were also analyzed. Quantitative qPCR experiments indicated that while *SlTGA9* showed reduced expression in the mutant in floral stages 8, 14 and 15, *SlTGA10* expression was strongly reduced in the mutant anthers from floral stage 8 and this low level persisted until floral stage16 (Figure **7B**, **C**). In Arabidopsis, ROXY1/ROXY2 glutaredoxins interact with TGA9/TGA10 transcription factors in the regulation of anther development (Murmu *et al*., 2010).

Globally, the expression analyses suggested that the absence of SlTPD1 and its downstream genetic network prevent the activation of the genes involved in the modulation of ROS levels, especially during early stages of anther development.

### ROS accumulation is lower in *Sltpd1* anthers than in the wild type at early developmental stages

The presence of reactive oxidative species was tested in the anthers of wild type and *Sltpd1* mutant plants. We analyzed and quantified the presence of superoxide anion (O2**^.-^**) and H_2_O_2_, considered the major ROS forms in plant cells (Huang *et al*., 2019), using 3-3’–diaminobenzidine (DAB) and nitroblue tetrazolium (NBT) staining, respectively. Quantification of NBT-staining of the anthers, a proxy for superoxide anion presence, detected the highest levels at floral stages 8 and 10 but no differences were observed between wild-type and mutant anthers (Figure **7A**). DAB quantification showed that in both wild-type and mutant anthers, the level of H_2_O_2_ is higher at floral stage 8 and then decreases progressively. Interestingly, at early stages (St8 and St10) the level of H_2_O_2_ was significantly lower in *Sltpd1* than in the wild type (Figure **7B**). These results suggest that a critical H_2_O_2_ threshold should be reached during early stages of anther development concurring with the meiotic stage.

In plants, the maintenance of ROS levels also relies on the action of non-enzymatic and enzymatic scavenging mechanisms. This last mechanism include enzymes such as, superoxide dismutase (SOD), catalase (CAT) and peroxidases (PRX) (Huang *et al*., 2019). To study the functionality of these enzymatic scavenging mechanism in the flowers of the mutant plants, we measured SOD and PRX activities. Compared to the wild type, SOD activity showed significant reduction in the mutant plants at floral stages 6 (premeiotic), 16 (pollen mitosis) and 20 (anthesis) (Figure **8C**). Remarkably, PRX activity was much reduced in *Sltpd1* mutant anthers (Figure **8D**) in agreement with the global downregulation of peroxidases shown in the RNA-seq experiment (Table **1**).

**Figure 8.**
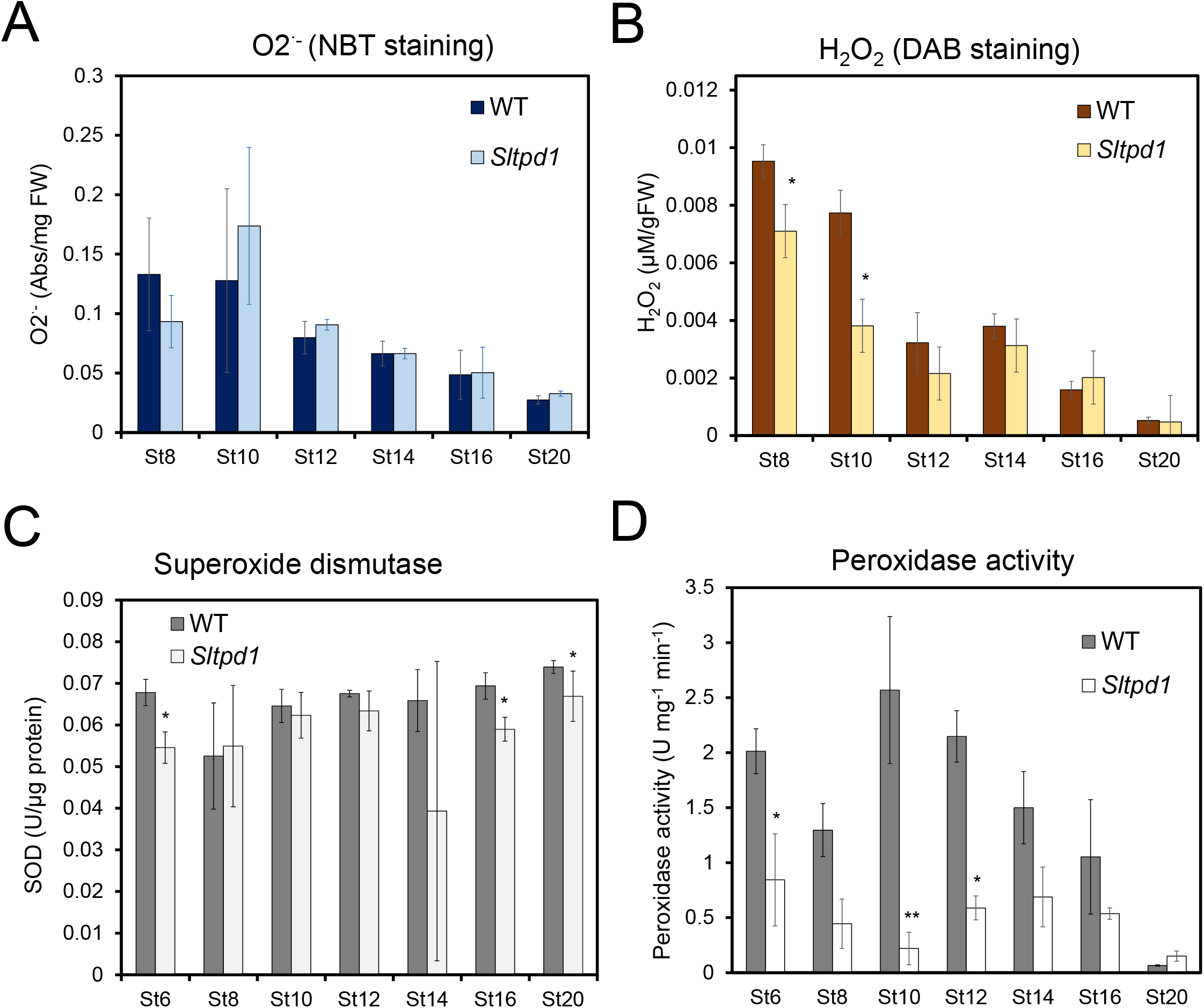
Redox homeostasis is altered in *Sltpd1* mutant anthers. Quantification of superoxide anion (O2·^-^) levels (A) and H_2_O_2_ levels (B) in wild-type and *Sltpd1* anthers at different developmental stages (n = 3 ± SD). Quantification of superoxide dismutase (C) and peroxidase (D) activity in wild-type and *Sltpd1* flowers at different developmental stages (St6-St20). Data correspond to 3 biological replicates ± SD. Statistical differences were inferred using a Mann-Whitney test. (*) = *p*<0.05, (**) = *p*<0.01.

## DISCUSSION

### *SlTPD1* is required for tapetum formation and pollen development in tomato

The stamens are the male reproductive organs of angiosperms and the place where the pollen is produced within the flower. The different tissues that compose anthers sequentially develop from the anther primordia suggesting that cell-to-cell communication is critical to coordinate growth and development (van der Linde and Walbot, 2019). The tapetum is the limiting tissue between the somatic and germinal cells and it is in a dynamic state during its short life period facilitating the pass of nutrients and molecules to the sporogenous cells and microspores (Pacini and Cresti, 1978).

In this study, we evince the pivotal role of the tapetal tissue during pollen development throughout the isolation and characterization of the *SlTPD1* gene. SlTPD1 shows homology with the TPD1 protein from Arabidopsis and, when transformed into the *tpd1* mutant, it was sufficient to complement the fertility defects. In addition, protein sequence alignments showed strong domain conservation also with the monocot proteins TDL1A/MIL2 and MAC1 from rice and maize. Therefore, SlTPD1 is the ortholog of the TPD1, MAC1 and TDL1A/MIL2 genes and the first gene ortholog identified in a fleshy fruit plant. On the other hand, although in tomato the putative receptor for SlTPD1 protein has not been yet identified, our data suggest the conservation of the receptor/ligand module also in tomato plants.

In Arabidopsis, mutant plants in either *EMS1* (TPD1 receptor) or *TPD1* genes share a phenotype, the lack of tapetum and the production of extra sporocytes at the expense of tapetal cells (Zh*ao et al*., 2002; Yang *et al*., 2003). We detected the expression of *SlTPD1* by *in situ* hybridization on the anther wall early in development until late stages where it appeared associated to the tapetum and microsporocytes. In this aspect, *SlTPD1* slightly differs from *TPD1* that is preferentially expressed in microsporocytes while *EMS* is predominantly expressed in tapetum (Zhao *et al*. 2002; Yang *et al*. 2003). In maize, MAC1 is expressed early in anther ontogeny where it suppresses archesporial cell proliferation, suggesting that cell position, rather than lineage regulates cell fate determination during anther development (Wang et al., 2012). This hypothesis is in agreement with the phenotype of *Sltpd1* mutants that showed defects in the shape and pattern of division of the archesporial cells. It has been shown that ectopic expression of *TPD1* activates cell division possibly by regulating the expression of cell-cycle genes (Huang et al., 2016a). Taken together, we propose a dual role for *SlTPD1* in the control of archesporial cell divisions and the determination of tapetal cell identity in tomato plants.

Most *TPD1* homologs are expressed in different tissues outside the anther including leaves, roots, seedlings (Yang *et al*., 2003; Hong *et al*., 2012a; Wang *et al*., 2012) and ovules (Yang et al., 2005; Wang et al., 2012). At present, a possible role of these proteins during vegetative development remains elusive. However, in monocots *TPD1* orthologs have been reported to control megaspore mother cell proliferation during ovule development (Sheridan et al., 1996; Zhao et al., 2008). Using *in situ* hybridization, the expression of *SlTPD1* was detected in anthers and the developing ovules. *Sltpd1* mutant plants did not show obvious defects in ovule development and flowers formed normal seeded fruits when pollinated with wild-type pollen. A peculiar and distinctive phenotype of the tomato *Sltpd1* mutants is the formation of seedless fruits (parthenocarpic). Parthenocarpy, the formation of fruits in the absence of pollination and fertilization, is often the consequence of the precocious activation of molecular events normally triggered by these processes (Molesini et al., 2020). Also, it could be achieved by external applications of different hormones or growth regulators (Vivian-Smith and Koltunow, 1999). In tomato plants, several reports suggest a role for developing stamens or male gametophytes in the repression of ovary growth (Medina et al., 2013; Hao et al., 2017; Rojas-Gracia et al., 2017; Okabe et al., 2019). Mutations in *SlTPD1* caused complete male sterility and the production of small parthenocarpic fruits. This phenotype could support this repressive effect exerted by male gametogenesis progression. Alternatively, the abnormal progression of male gametogenesis could result in the production of signaling molecules that indirectly activate premature ovary growth. In this regard, antisense plants targeting *SlRBOHB*/*SlWfi1* a tomato gene involved in the generation of ROS, show several developmental defects including parthenocarpic fruit development (Sagi et al., 2004).

### *SlTPD1* and the control of redox homeostasis during pollen development

Overlapping with the genetic network controlling anther development, additional factors and signalling molecules participate in the communication between the somatic and sporogenous tissues. These factors include hormones, secreted proteins, miRNAs and cellular redox state (Dukowic-Schulze and van der Linde, 2021). For instance, the analysis of gibberellin (GA) deficient mutants suggest that the primary site of hormone action are tapetal cells and low GA levels have an indirect effect on the formation of functional pollen grains (Aya *et al*., 2009). Several lines of evidence support that cellular redox state is an important morphogenetic factor controlling cell differentiation and proliferation during anther development (Yu and Zhang, 2019).

Interestingly, while high concentration of reactive oxygen species (ROS) cause irreversible DNA damage and cell death, at low levels ROS act as signalling molecules regulating cell division and cell fate (Kelliher & Walbot, 2012; Yang *et al*., 2018). The results presented in this study show that the absence of tapetal tissue in *Sltpd1* mutants have a huge impact in the transcription of genes involved in redox homeostasis in the anther at early stages. Moreover, a reduction in ROS levels seem to be associated with the failure of pollen mother cells to progress into meiosis. In agreement with this observation, pioneering work in maize showed that hypoxia triggers meiotic fate acquisition acting as a positional cue for germ cell production (Kelliher and Walbot, 2012).

Besides the production of ROS as an end product of several metabolic processes, a specific enzymatic machinery is in charge of maintaining redox homeostasis in plants. ROS production relays on *RBOH* genes also known as NADPH oxidases, which catalyze the generation of superoxide radicals. Enzymatic scavenging mechanism involve SOD, CAT and peroxidases, although peroxidases can act as both ROS-generating and ROS-processing components (Mittler, 2017). Cellular changes of ROS levels can act as a signal to regulate differentiation and morphogenesis during reproductive development. In Arabidopsis, *RBOHE* is specifically expressed in the tapetum and the genetic interference with the temporal ROS pattern resulted in altered tapetal PCD and male sterility (Xie *et al*., 2014). In addition, PRX9 and PRX40 are extensin peroxidases specifically expressed in the tapetum that act as scavenging molecules contributing to tapetal cell wall integrity (Jacobowitz et al., 2019). ROS signalling include glutaredoxins (GRXs) that act as sensors of the redox status, altering signal transduction pathways that result in biological responses (Song et al., 2002). Studies in Arabidopsis, rice and maize highlighted the importance of GRXs in the formation of the anther and the differentiation of microsporocytes (Xing and Zachgo, 2008; Hong et al., 2012b; Kelliher and Walbot, 2012). In rice, a mutation in the anther-specific glutaredoxin *MICROSPORELESS1* (*MIL1*) prevent the completion of the meiosis during male gametogenesis. *MIL1* encodes a CC-type glutaredoxin that specifically interact with TGA transcription factors (Hong et al., 2012b). In Arabidopsis, *ROXY1* and *ROXY2* encode also CC-type glutaredoxins and are required for the formation of the adaxial anther lobe possibly with other GRXs or redox regulators. ROXY1 and ROXY2 proteins are able to interact with the TGA transcription factors TGA9 and TGA10 in tobacco leaves (Xing and Zachgo, 2008). These authors suggest that this interaction results in the modification of TGA9/10 and its activation as a transcriptional factor (Murmu et al., 2010). A similar genetic network to the one described in rice and Arabidopsis should operate in tomato anthers where ROS produced in the tapetal cells orchestrate anther wall development and pollen mother cells progression into meiosis. Using a tomato mutant lacking the tapetum, we identified several elements of this network that were included in the proposed working model (Figure **9**). ROS produced by RBOHs (*SlRbohB* and *SlRbohE*) and peroxidases results in the accumulation of H_2_O_2_ in tapetal tissue. Glutaredoxins, including *SlGRXC9*, could target TGA transcription factors (*SlTGA9* and *SlTGA10*) for activation, regulating then the expression of a set of genes required for further stages of pollen and anther development. This genetic network is severely affected by the absence of *SlTPD1* and the concomitant loss of the tapetal tissues, at early stages of anther formation.

**Figure 9.**
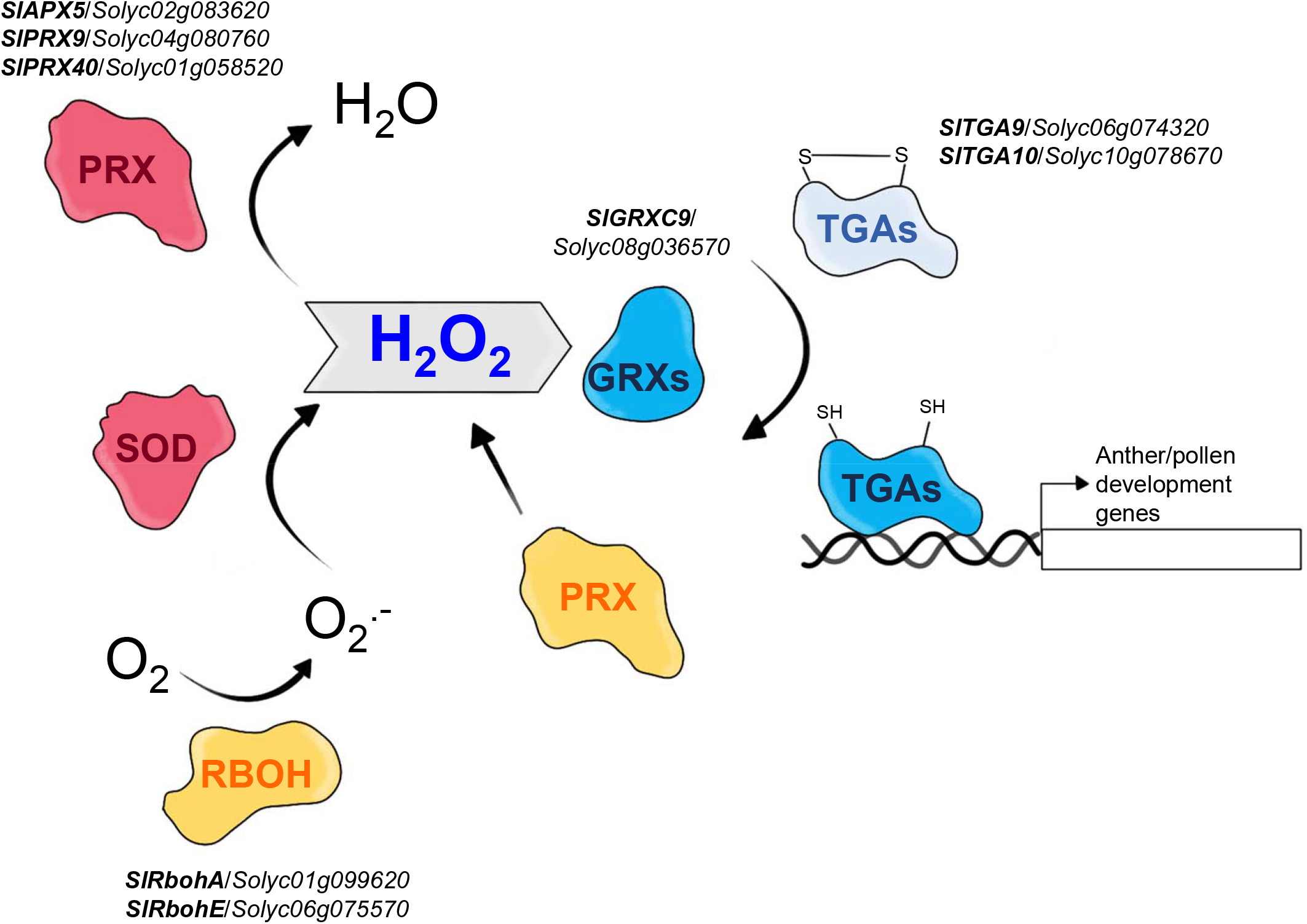
Working model summarizing the genetic elements of the redox network affected by the absence of *SlTPD1* and the concomitant tapetum loss, at early stages of tomato anther development. Enzymatic ROS accumulation (Orange) is attenuated by ROS scavenging mechanism (Pink). Changes in ROS levels activate signalling pathways (Blue) that result in the induction of genes involved in anther/pollen development. APX: ascorbate peroxidase; GRXs: glutaredoxins; PRX: peroxidases; RBOH: Respiratory burst oxidase homolog; SOD: superoxide dismutase; TGAs: TGA transcription factors.

In the present study, we identified and characterized the tomato gene *SlTPD1* that has a central role in pollen formation. Using CRISPR/Cas9 technology, we generated male sterile tomato plants that specifically lack tapetal tissue to gain insight into the genetic network and molecular signals regulating pollen formation in this species. Based on the obtained information, a working model is proposed highlighting the role of ROS production and signalling during early stages of pollen development in tomato plants.

## MATERIALS AND METHODS

### Plant material and growth conditions

Tomato (*Solanum lycopersicum* L.) seeds from cultivar Moneymaker were grown in a greenhouse in pots with a 2:1:1 mixture of peat:vermiculite:perlite with a temperature regime of 25-30°C day and 18-20 °C night. Natural light was supplemented with lamps to obtain a 16h light/8 h night photoperiod. Flower samples were collected at different developmental stages according to bud size (Brukhin *et al*., 2003). In this study, 8 floral stages were analyzed that correspond to the following sizes: St6 (0.3 cm), St8 (0.4 cm; meiotic), St10 (0.5 cm; tetrad of microspores), St12 (0.6 cm), St14 (0.7 cm), St16 (0.8 cm; pollen mitosis), St18 (0.9 cm) and St20 (1 cm; anthesis). For the characterization of tomato fruits, size and weight of at least thirty fruits from the different genotypes were analyzed when fully ripe.

*Arabidopsis thaliana* Columbia (Col) plants were used as the wild-type genotype. The *tpd1* mutant corresponds to the T-DNA insertion line N843482 (SAIL_1174_B09) obtained from Nottingham Arabidopsis Stock Centre (NASC). The line was genotyped using the T-DNA specific primer LBb1 and the gene specific primer pair TPD1-LP1/TPD1-LP2 that amplifies 427 bp from the *TPD1* gene (Supplementary Table **S2**). Arabidopsis plants were grown in seed pots in a growth chamber with a 2:1:1 mixture of peat:vermiculite:peat at 21°C under long day conditions (16h light/8 h dark) and irrigated with Hoagland’s solution.

### Phylogenetic analysis

The phylogenetic tree was inferred by the neighbor-joining method using Poisson-corrected amino acid distances. A total of 1000 bootstrap pseudo-replicates were used to estimate reliability of internal nodes. Tree inference was performed using MEGA version 6 (Tamura *et al*., 2013). The dataset comprised 18 *TPD1*-like genes from different plant species, obtained from GenBank database (Table **S1**).

### Plasmid assembly

Construct for the complementation of the Arabidopsis *tpd1-2* mutant: A fragment of 2.7 kb from the *TPD1* promoter (5’ region of the gene) was fused to the coding sequence of the tomato *SlTPD1* gene. First, both fragments were independently amplified by PCR and cloned into the intermediate vectors pENTRY 5’ TOPO and pCR8 (Invitrogen), respectively. Second, a multisite gateway recombination reaction was performed to introduce both sequences into the binary destination vector pK7m24GW,3 (https://gateway.psb.ugent.be) to obtain the final construct *pAtTPD1::SlTPD1*.

Design of gRNA and CRISPR/Cas9 construct for *SlTPD1* gene editing: For the design of optimal gRNAs, the target site was selected using the Breaking-Cas design tool (Oliveros *et al*., 2016). This tool is freely available *on line* (https://bioinfogp.cnb.csic.es/tools/breakingcas). CRISPR/Cas9 plasmid assembly was performed using the Golden Braid (GB) modular framework and tools (http://www.gbcloning.upv.es). First, a single gRNA sequence was obtained by annealing of complementary primers and then assembled with GB1001 (U626 promoter) and GB0645 (scaffold RNA) parts into the destination vector pDGB3α1. In successive multipartite GB reaction, this first module was assembled together with the GB0639 and GB0226 parts (containing hCas9 and *nptII* transcriptional units, respectively) into the final destination vector. The final construct was then transformed into *Agrobacterium tumefaciens* strain LBA4404. The primers used are listened in supplementary Table **S2**.

### Plant transformation

Arabidopsis transgenic plants were obtained using the floral dip method (Clough and Bent, 1998). Briefly, plants were grown under long day conditions until flower transition occurs and then the main stem was removed to allow growing of secondary meristems. *Agrobacterium* inoculation (C58C1 strain carrying construct of interest) was performed by immersion of the shoots (2-5 cm length) in a suspension containing 5% sucrose and 0.05% Silwet L-77. Transformant plants were selected in the presence of kanamycin and transferred to soil for further analyses.

Tomato transformants were obtained by *in vitro* co-cultivation of the *Agrobacterium* strain LBA4404 (carrying the binary vector of interest) and cotyledon explants (Ellul *et al*., 2003). Transformants were selected in the presence of kanamycin and after rooting, transferred to the greenhouse.

### Genotyping of CRISPR/Cas9 edited plants

Genomic DNA was extracted from young leaves or unopened flower buds. A 530 bp fragment from the *SlTPD1* genomic region flanking the targeted region was amplified using oligos SlTPD1G For and SlTPD1G Rev, purified and sequenced. T0 plants with percentages of edition over 80% were selected using the *on line* tool TiDE (http://shinyapps.datacurators.nl/tide/) (Brinkman et al 2014). We then used the *on line* software ICE v2 CRISPR analysis tool (https://ice.synthego.com/) to identify the number and type of edition for each plant.

For the genotyping of stable and Cas9-free edited plants, PCR-based molecular markers were designed. We used Cleaved Amplified Polymorphic Sequences (CAPS) markers (Konieczny and Ausubel, 1993) that detect polymorphisms that occur in restriction sites. The deletions present in the *Sltpd1^del2^* and *Sltpd1^del5^* allele, generated new restriction sites for *Bse*GI and *Nco*I enzymes, respectively. Using SlTPD1G For and SlTPD1G Rev oligos (Table **S2**) a 530bp fragment was obtained from genomic DNA. *Bse*GI generated two fragments of 308 bp and 220 bp in the *Sltpd1^del2^* allele and *Nco*I generated two fragments of 299 bp and 226 bp in the *Sltpd1^del5^* allele. Neither of the enzymes cut the wild type fragment.

### Subcellular localization of SlTPD1

The coding sequence of *SlTPD1* was cloned via Gateway LR reaction into the pEarleyGate101 vector (YFP fluorescent tag-containing)(Earl*ey et al*., 2006) to generate the expression vector SlTPD1-YFP. The vector was transformed into *Agrobacterium tumefaciens* strain C58 and used to agroinfiltrate 4-weeks-old *Nicotiana benthamiana* leaves. After 48 hours of the infiltration, the localization of the fluorescence fusion protein was determined on leaf disks by confocal scanning microscopy (LSM 780, Zeiss, Jena, Germany). A 35S:GFP construct was used as a control.

### Expression analyses by quantitative real-time PCR (qPCR)

Total RNA was extracted from frozen tissue using the E.Z.N.A. Plant RNA Kit (Omega BioTek). RNA was treated with DNAse I (Thermo Fisher Scientific) to remove genomic DNA and quantified in a NanoDrop ND-1000 Spectophotometer (Thermo Fisher Scientific). For first-strand cDNA synthesis, one microgram of DNase-treated RNA was used for reverse transcription using a PrimerScript RT reagent kit (Takara) and a mix of oligo poli-dT and random hexamers. The resulting cDNA was used for quantitative RT-PCR with the MasterMix qPCR ROX PyroTaq EvaGreen 5x (CmB) and the reaction was run on a QuantStudio 3 (Applied Biosystems). Relative expression levels were calculated by normalizing to the reference genes *ACT* (Arabidopsis experiments) or *SlActin8* (tomato experiments) and using the ΔΔCt method. All primers showed amplification efficiencies between 90 and 110%. The primers used are listed in the supplementary Table **S2**.

### RNA *in situ* hybridization in tomato flowers

Fresh floral samples were fixed in FAE (4% formaldehyde, 5% acetic acid, 50% ethanol) overnight at 4°C, and afterwards, stored in 70% ethanol. Samples were embedded in paraffin using an automated tissue processor (Leica TP1020).

To generate gene specific probes cDNA fragments were cloned under T7/SP6 promoter sequences. For *SlTPD1* a 284 bp DNA fragment from the 5’ coding region was amplified by PCR using cDNA from flowers and cloned into the pGEM-T Easy vector (Promega). For *TomA5B* and *SlSDS* genes, we used cDNA fragment of 442 bp and 440 bp respectively. Digoxigenin-labelled probes were transcribed *in vitro* with T7 or SP6 RNA polymerases. RNA was hybridized *in situ* (Huijser et al., 1992; Gómez-Mena and Roque, 2018) in paraffin-embedded sections (8µm) and color was detected with 5-bromo-4-chloroindol-3-yl phosphate/nitrateblue tetrazolium (BCIP/NBT) (Roche).

### Histological techniques

For histological studies, tissue was fixed in FAE overnight at 4 °C and stored in 70 % ethanol. Samples were embedded in acrylic resin (Technovit 7100; Kulzer) according to the manufacturer’s instructions. For histological analysis, resin sections were stained with 0.05 % toluidine blue in 0.1 M 6.8 pH phosphate buffer (O’Brien *et al*., 1964) and visualized in a Leica DM 5000B microscope (Leica Microsystems) under bright field.

### Aniline blue staining in cryosections

For assays in which fresh tissue was needed, samples were fixed in NEG-50 (Richard Alan Scientific), rapidly frozen in liquid nitrogen, and cut into 16 µm sections using a cryostat (Microm HM 520). Cryosections were stained for 10 minutes in the darkness with 0.5 % aniline blue in 0.07 mM sodium phosphate buffer, and visualized in a Leica DM 5000B microscope (Leica Microsystems).

### Pollen viability essay

Alexander’s staining was carried out as previously described (Peterson *et al*., 2010) with 2 minutes of incubation at 50 °C on a hot plate. For pollen viability, pollen was released from the anthers by squeezing, stained and counted. Samples were visualized in a Leica DM 5000B (Leica Microsystems) microscope under bright field. For each sample, thirty anthers from five different flowers were used.

### Histochemical localization and quantification of hydrogen peroxide (H_2_O_2_) and superoxide radical (O2^·-^)

Hydrogen peroxide localization was performed in anthers obtained from flowers in different developmental stages. Immediately after dissection, anthers were submerged in a 1 mg ml^-1^ DAB-HCl (pH 3.8) solution for 16 hours under light conditions (Unger *et al*., 2005), then cleared in 80 % ethanol for 20 minutes and observed in a binocular microscope (Leica Microsystems). Hydrogen peroxide levels were quantified following a similar method. After staining in DAB-HCl and clearing with ethanol, anthers were pulverized in liquid nitrogen, dissolved in 0.2 M HClO4 and centrifuged at 12000g for 10 minutes. The absorbance of the supernatant was quantified at 450 nm. H_2_O_2_ concentrations were obtained through a standard curve for known hydrogen peroxide concentrations diluted with 0.2 M HClO4-DAB (Kotchoni et al., 2006).

Superoxide radical was measured as formazan formation over time from tetrazolium blue. Flowers from different developmental stages were weighted, submerged in 50 mM potassium phosphate buffer (pH 7.8) containing 0.1% NBT and 10 mM sodium azide, left to stain for 2 hours and cleared in 70% ethanol. After staining, tissue was rapidly frozen in liquid nitrogen and ground. Formazan was selectively extracted using 200 µl of DMSO and absorbance was measured at 550 nm.

### Peroxidase (PRX) and superoxide dismutase (SOD) activity

Flowers at different developmental stages were collected and frozen in liquid nitrogen. Frozen tissue was ground and homogenized in extraction buffer (0.1M Tris pH 7.0, 0.1% ascorbic acid, 0.1% L-cysteine, 0.5M sucrose and 10mg/ml PVP) and centrifuged at 4°C for 15 minutes, saving the supernatant. Total protein was quantified using the Bradford method (Bradford, 1976). Briefly, 10 µl of crude extract were added to a tube containing 1 ml of Bradford solution (0.01% Coomasie Brilliant Blue G-250, 4.7% ethanol, 8.5% phosphoric acid) and mixed. After two minutes, the absorbance was measured at 595 nm. A standard curve was generated using known concentrations of BSA.

For SOD activity, 25 mg of protein from the crude extract were added to 1ml of SOD buffer (50mM PBS pH7.6, 0.01mM EDTA, 50mM sodium carbonate, 12mM L-methionine, 10 µM riboflavin, 50 µM NBT) and incubated at room temperature under light conditions for 10 minutes. Absorbance was measured at 550 nm and SOD buffer without extract was used as a negative control. SOD activity was quantified as the amount of enzyme required to inhibit 50% of the photoreduction of NBT.

For PRX activity, 25 to 75 mg of protein from the crude extract were added to 1 ml of PRX buffer (0.85 mM hydrogen peroxide in HEPES pH7.0, 0.125M 4-aminoantipyrene, 8.1 mg/ml phenol) and the change in absorbance was measured for 2 minutes at 510 nm. A standard curve was generated using known concentrations of horseradish peroxidase.

### RNA-Seq analyses

Total RNA was extracted from stage 8 stamens from the wild type and *Sltpd1* plants. Frozen tissue using a NucleoSpin RNA Plant kit (Mascheny-Nagel) and measured in a NanoDrop ND-1000 Spectophotometer (Thermo Fisher Scientific). The RNA quality was assessed based on the RNA integrity number (RIN) using Bioanalyzer 2100 (Agilent) and samples with RIN>8 were selected for the experiment. RNA sequencing was performed using the BGISEQ Technology platform at BGI (China). A total of three biological replicates were used for each sample set. GO enrichment, KEGG enrichment and statistical analysis were done through the Dr. Tom platform (BGI, China).

### Statistical analysis

IBM SPSS Statistics v.27 was used for statistical analysis. For each data set, a Shapiro-Wilk normality test was run. For normally distributed data, a Student-t test was used for pairwise comparison. Non-normally distributed data were analyzed with a Mann-Whitney test.

## SUPPLEMENTAL DATA

Figure S1. Expression pattern of two tomato *TPD1* gene homologs analyzed by quantitative RT-PCR in leaves and floral buds at different developmental stages.

Figure S2. *In silico* analyses of Arabidopsis and tomato *TPD1* gene homologs. Figure S3. Expression of *SlTPD1* in the ovary detected using *in situ* hybridization.

Figure S4. Characterization of CRISP/Cas9-mediated *SlTPD1* edited tomato plants. Table S1. Accession numbers of *TPD1*-like gene sequences from different plant species used for the phylogenetic analysis.

Table S2. Oligonucleotides used in this study.

Table S3. List of differentially expressed genes (DEGs) between wild-type and *Sltpd1* mutant anthers from floral stage 8.

## ACKNOWLEDGMENTS

This work was supported by grant RTI2018-094280-B-I00 funded by MCIN/AEI/ 10.13039/501100011033 and by FEDER “A way of making Europe”. We thank Aureliano Bombarely for his help in the conversion of gene IDs into Solyc identifiers, Diego Orzáez for providing GoldenBraid parts and Maricruz Rochina for expert technical assistance during the project.

